# PD-1 Targeted Antibody Discovery Using AI Protein Diffusion

**DOI:** 10.1101/2024.01.18.576323

**Authors:** Colby T. Ford

## Abstract

The programmed cell death protein 1 (PD-1, CD279) is an important therapeutic target in many oncological diseases. This checkpoint protein inhibits T lymphocytes from attacking other cells in the body and thus blocking it improves the clearance of tumor cells by the immune system. While there are already multiple FDA-approved anti-PD-1 antibodies, including nivolumab (*Opdivo*® from Bristol-Myers Squibb) and pembrolizumab (*Keytruda*® from Merck), there are ongoing efforts to discover new and improved checkpoint inhibitor therapeutics. In this study, we present multiple anti-PD-1 antibody fragments that were derived computationally using protein diffusion and evaluated through our scalable, *in silico* pipeline. Here we present nine synthetic Fv structures that are suitable for further empirical testing of their anti-PD-1 activity due to desirable predicted binding performance.

## Introduction

The field of cancer immunotherapy has witnessed unprecedented advancements over the past few decades, reshaping the landscape of oncological treatment strategies and extending the lives of those with disease. Among the notable breakthroughs is the identification and targeting of programmed cell death protein 1 (PD-1), a crucial immune checkpoint receptor that plays a pivotal role in modulating T-cell responses. PD-1 belongs to the CD28 superfamily and is predominantly expressed on activated T cells (1). PD-L1 and PD-L2, the ligands of PD-1, are highly expressed on multiple types of cancer cells and thus plays an important role in immune evasion (2, 3).

PD-1 emerged as a key focus in cancer research due to its ability to suppress the immune system and facilitate immune evasion by tumor cells. This discovery has strengthened the development need of innovative therapeutic interventions designed to support the immune system’s full potential against cancer (4).

Pembrolizumab (marketed as Keytruda® by Merck) and nivolumab (marketed as Opdivo® by Bristol Myers Squibb), two of the first monoclonal antibodies targeting PD-1, first gained FDA approval for the treatment of melanoma in 2014 (5, 6). Both, along with other antibodies, have since been extended approval to various other malignancies including non-small cell lung cancer, head and neck squamous cell carcinoma, and Hodgkin’s lymphoma, colorectal cancer, renal cell carcinoma, and others (7–11).

Pembrolizumab and other similar antibodies have demonstrated remarkable efficacy across a variety of cancers, offering durable responses and improved survival rates (12, 13). Despite the success of PD-1-targeted therapies, challenges persist, including treatment resistance (14), variability in patient response (15, 16), and the need for personalized approaches (17). Herein lies the motivation for exploring novel strategies in the design and optimization of PD-1-targeting antibodies.

This study explores the use of artificial intelligence (AI) in the design of antibodies through protein diffusion. Protein diffusion is a new technique that uses deep learning-based models to generate amino acids sequences. This can be performed “unconditionally”, where the AI model generates a random sequence of a desired length, or “conditionally”, where there system has some reference data and will produce proteins that mimic a given input protein class. We can then fold the diffused amino acid sequences to produce protein structure files to be used in subsequent analyses.

Using a large corpus of existing anti-PD-1 antibodies to conditionally guide the AI system, we have generated 9 antibody candidates and assessed their viability as compared to other therapeutics through *in silico* protein-protein docking.

By leveraging the power of protein diffusion, we aim to contribute to the evolving landscape of cancer immunotherapy, offering insights that may lead to the development of next-generation PD-1-targeted antibodies. This approach holds promise for addressing current development bottlenecks and advancing the field towards more effective and personalized treatments for cancer patients. In the following sections, we delve into the methodology and potential implications of utilizing protein diffusion in the design of PD-1-targeting antibodies, aiming to contribute to the ongoing efforts in the pursuit of precision oncology.

## Methods

Heavy and light chain sequences of 33 PD-1 targeting antibodies (Fv region only) were retrieved from the Therapeutic Structural Antibody Database (Thera-SAbDab) (18) on November 11, 2023. The sets of heavy and light chain sequences were each aligned using Muscle v3.8.425 (19), pro-ducing .fasta files of the alignments. These were then converted to the .a3m format.

EvoDiff, a suite of protein generation models from Microsoft Research (20), was used for diffusion of new antibody structures that target PD-1. This generative framework uses an input of aligned amino acid sequences for conditional diffusion where the diffusion process is “evolutionarily-guided” through predictions based on the input set.

Conditional diffusion was carried out using the .a3m alignment files with EvoDiff’s `MSA_OA_DM_MAXSUB` model using the `generate_query_oadm_msa_simple()` function. Three heavy chain Fv sequences and three light chain Fv sequences were diffused through this process, as shown in Table 1. Then, these were combined to generate 9 antibody candidates, as listed in Table 2.

**Table 1.**
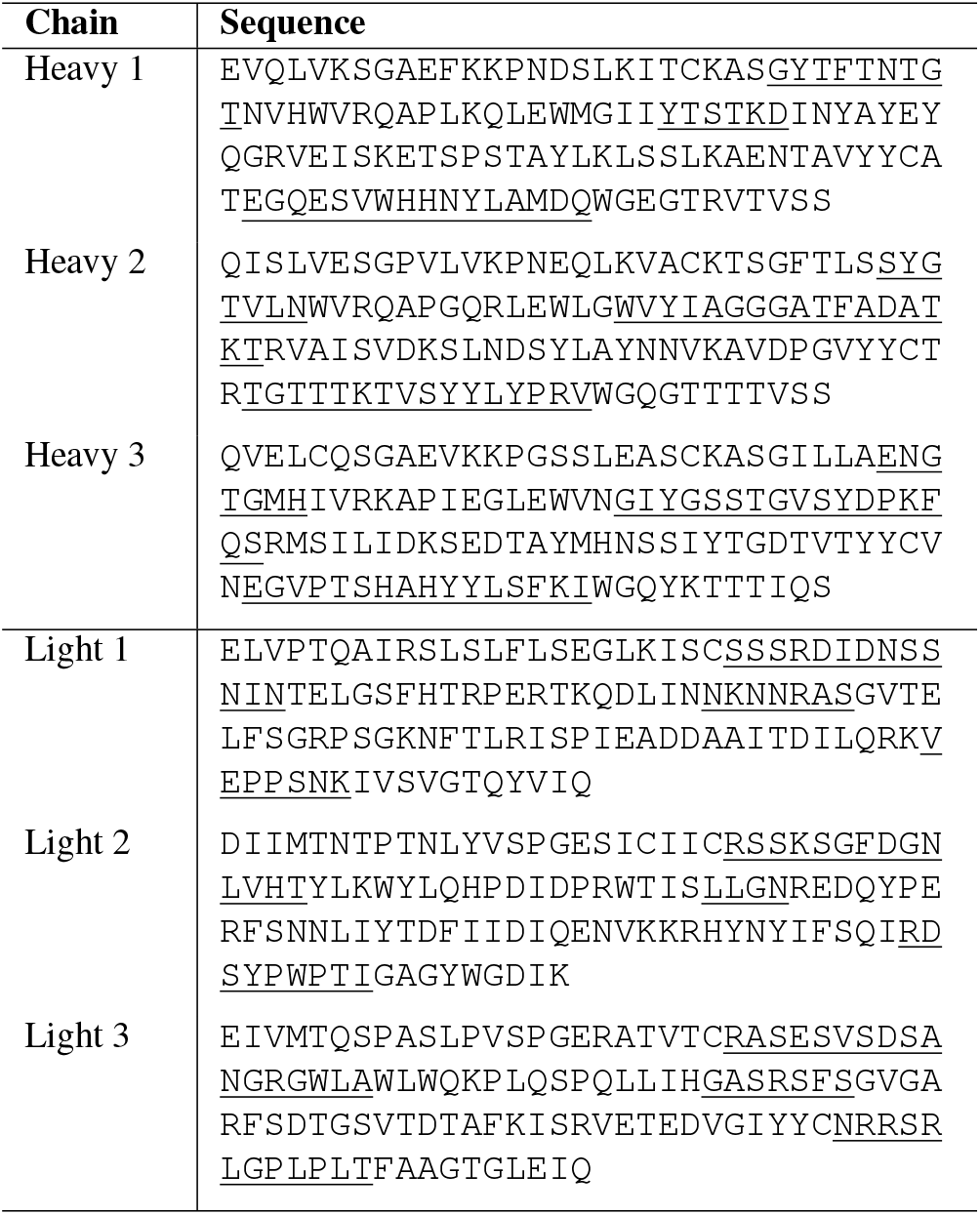
Conditionally-diffused sequences of the PD-1-targeting antibody candidate heavy and light chains. CDR loop residues are <u>underlined</u>.

**Table 2.** Heavy and light chain assignments of the PD-1-targeting antibody candidates.

| Antibody ID | Heavy Chain | Light Chain |
| --- | --- | --- |
| TUPPD1-001 | 1 | 1 |
| TUPPD1-002 | 1 | 2 |
| TUPPD1-003 | 1 | 3 |
| TUPPD1-004 | 2 | 1 |
| TUPPD1-005 | 2 | 2 |
| TUPPD1-006 | 2 | 3 |
| TUPPD1-007 | 3 | 1 |
| TUPPD1-008 | 3 | 2 |
| TUPPD1-009 | 3 | 3 |

### Sequence Analysis

The diffused sequences were analyzed through abYsis, an antibody analysis tool (21), to highlight the parts of the Fv regions of the antibodies, especially the CDR loop residues. This tool also highlights any residues that may be unusual (occurring in <1% of sequences) as compared against a corpus of 100,357 antibodies from *Homo sapiens*. In addition, the tool identifies potential sites for oxidation, phosphorylation, glycosylation, etc.

### Structure Prediction

Using ColabFold v1.5.5 (22) Batch, which utilizes AlphaFold2 Multimer (23, 24) with MMseqs2 (25), the Fv structures for the diffused sequences in Table 1 along with the 33 Thera-SAbDab references were predicted. The side chains of these predicted structures were relaxed using the OpenMM/Amber method (26) in ColabFold.

This process generates a .pdb file for each of the Fv candidate structures. The L chain in each of these .pdb files was renumbered using PyMol v2.4.1 (27) to avoid duplicative residue numbering with the H chain, which is a requirement for the docking process.

### Protein Docking

HADDOCK v2.4 is a biomolecular modeling software that was used to dock the structures in this study (28). This tool provides docking predictions for provided structures using an information-driven flexible docking approach.

For this study, we utilized a Docker containerized version of HADDOCK^1,2^, which contains all of the software dependencies to allow HADDOCK to run more readily in an high-performance computing (HPC) environment. HADDOCK was run on 36 physical cores in the University of North Carolina at Charlotte HPC cluster.

To prepare for protein docking, active residues must be determined for both the antibody and antigen in each experiment. For the PD-1 antigen, the active residues were determined to be those as the interfacing site between PD-1 and PD-L1. Namely, ASN66, THR76, LYS78, and surrounding residues. This is the common binding site for many of the therapeutic antibodies, including pembrolizumab.

As for the Fv structures, active residues were deemed to be the residues in the CDR loops. Residues in the CDR loops were programmatically detected using the ANARCI system (29). This process returns the residues numbers, based on the Chothia numbering system, for the CDR1, CDR2, and CDR3 loops.

The HADDOCK experiment files were programmatically generated using custom Python logic, which created directories for each of the antibodies and placed the required files in each directory. Then, each experiment was submitted to the HPC cluster to be run in parallel across the distributed compute nodes. To complete the 42 docking experiments, this took approximately 3 hours on ten 36-core nodes. This scalable docking process closely follows methods reported in Tomezsko and Ford et al. 2023 (30) and the published antibody docking protocol from the Bonvin Lab at Utrecht University (28, 31, 32).

### Protein Complex Evaluation

Upon completion of the HADDOCK iterations, the relevant outputs were collected, including the metrics and .pdb complex files from the water refinement stage of the docking process. Using the reported van der Waals (VDW) energy metric, the metrics of the top performing cluster were recorded. Then, the “best” .pdb file, in terms of lowest VDW energy, from the top performing cluster was selected as the representative structure for subsequent analyses, visualizations, and comparisons.

Also, PRODIGY, a tool to predict the binding affinity of protein-protein complexes, was used on each “best” structure for each complex in this study (33). The predicted binding affinities are reported as Gibbs energy, shown as ΔG (in Kcal/mol units).

Protein structures and complexes resulting from the diffusion/Thera-SAbDab procurement process and the HADDOCK docking processes, respectively, were visualized using PyMOL v2.4.1 (27). PyMOL was also used to help select active residues at the PD-1/PD-L1 interface from PDB: 3BIK and to evaluate the antibody active residues as selected by abYsis and manual selection. Then, interfacing residues were detected (polar contacts within 3.0Å) between the PD-1 and Fv complexes that may indicate potential inhibition activity.

### Molecular Dynamics

Molecular dynamics analyses were performed using OpenMM on the predicted complexes of TUPPD1-001, TUPPD1-002, TUPPD1-009, pembrolizumab, and nivolumab (26). The complex PDBs were prepared using the OpenMM PDBFixer tools and then solvated with a an ionic strength of 150mM and a pH of 7.4. Protein topologies were generated with the CHARMM36 force field in water (34). For the evaluation of Coulombic interaction, the particle mesh Ewald (PME) setting was applied to the force field system (35). The pressure and temperature simulations were set using the MonteCarloBarostat method (pressure = 1 bar and temperature = 300 K) and Langevin dynamics (*γ*_*Lang*_ = 1/ps), respectively. The simulations were then run for 100,000 steps where 1,000 steps equaled 4 picoseconds.

The results of these analyses, including the solvated and energy minimized PDB files and equilibrated metrics, are provided in the Supplementary Materials in the GitHub repository.

## Results

Through protein diffusion, 9 antibody candidates were generated that bind similarly to other existing therapeutic antibodies. As the binding site on PD-1 was constrained to be the normal interface between PD-1 and PD-L1, which is also the binding site for other commercially available antibodies like pembrolizumab and nivolumab, the docking results show consistent interactions in this area. However, the binding orientation and angles of these diffused antibodies against PD-1 vary. These docking results are shown in Figure 1.

**Fig. 1.**
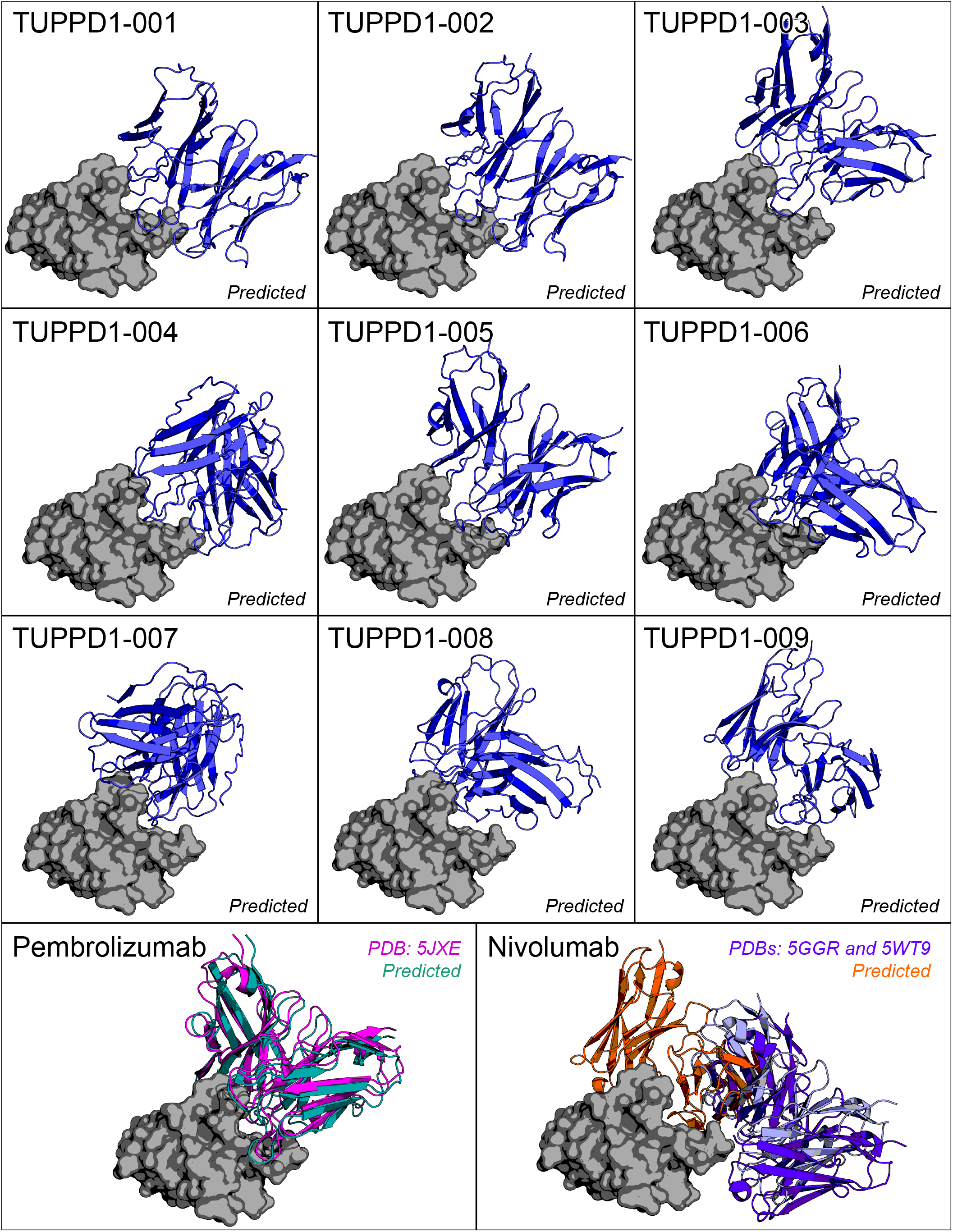
Predicted docking complexes of nine conditionally-diffused antibodies. The grey surface protein is PD-1 and the cartoon structures are the Fv portions of the antibodies. The docking location of pembrolizumab is shown in teal (predicted) and pink (actual, PDB: 5JXE) and the contested docking location of nivolumab is shown in orange (predicted) and light purple/purple (actual, PDBs: 5GGR and 5WT9, respectively).

As shown in Figure 2, some of the 9 diffused candidates bound to PD-1 with similar affinities, though not in all cases nor across all metrics.

**Fig. 2.**
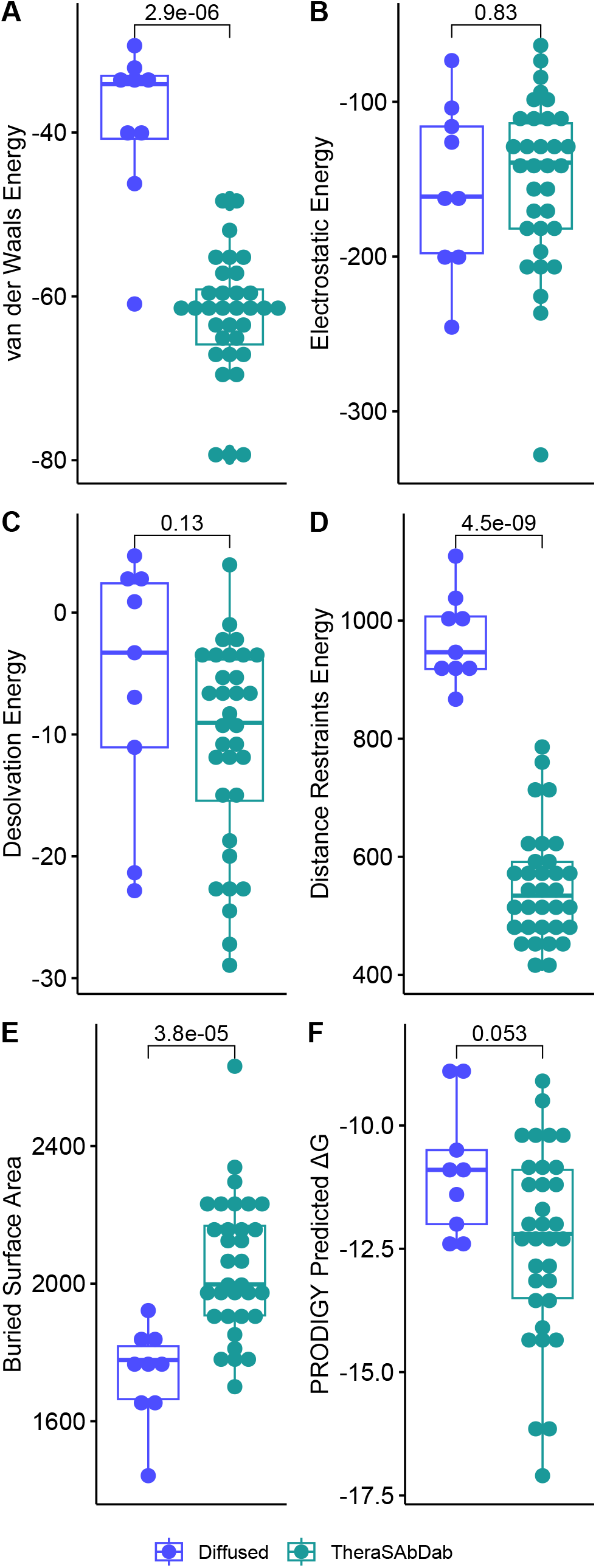
Comparison of the docking metrics between existing and diffused antibodies. Pairwise comparisons are shown as p-values from the Wilcoxon signed-rank test. For all metrics except buried surface area, lower is likely indicative of better binding.

### Structure Analysis

For the 9 antibody candidates that were generated through protein diffusion, all folded into tertiary structures that resemble normal immunoglobulin G structures, including the other 33 antibodies procured from Thera-SAbDab. The diffused antibody chains were all classified as heavy chain subgroup I and kappa light chain subgroup IV through abYsis.

However, there are certain features of the diffused heavy and light chains that are unnatural when compared to biologically-derived structures. For example, abYsis identified numerous residues in all of diffused chains that are unusual, meaning residues were picked in the diffusion process in certain positions that are uncommon as compared to naturally-derived human antibodies. Also, in light chains 1 and 2 there are secondary structure differences such as the extension or reduction, respectively, of the anti-parallel beta sheet between complementarity-determining region 3 (CDR3) and frame region 4 (FR4) as compared to other light chains in the reference set. It is unknown as to the effect of these secondary structure differences, though the confidence in the protein folding prediction remains high in all areas except for the CDR loops, which is expected. Of note, candidates TUPPD1-001, TUPPD1-002, and TUPPD1-009 showed the most promise as their binding metrics were the strongest across multiple biochemical features. Each of these candidates exhibit favorable predicted van der Waals, electrostatic, and Gibbs energies as compared to the larger Thera-SAbDab reference set. See Table 3.

**Table 3.** Comparison of the three top performing diffused antibodies against nivolumab and pembrolizumab, along with group averages.

| Antibody ID | van der Waals Energy | Electrostatic Energy | PRODIGY Predicted $\Delta G$ |
| --- | --- | --- | --- |
| TUPPD1-001 | -60.93 | -161.23 | -10.90 |
| TUPPD1-002 | -46.24 | -73.40 | -11.40 |
| TUPPD1-009 | -40.76 | -202.88 | -12.50 |
| *Diffused Average* | *-38.78* | *-154.52* | *-10.92* |
| Nivolumab | -78.67 | -204.76 | -14.40 |
| Pembrolizumab | -62.22 | -328.10 | -11.70 |
| *Thera-SAbDab Average* | *-62.32* | *-150.75* | *-12.38* |

Furthermore, molecular dynamics simulations show that the predicted HADDOCK output structures were in a nearly optimal energy minimized state. For example, when aligning the predicted complexes from HADDOCK versus the energy minimized versions from OpenMM, the RMSD of the alignments were very low, suggesting considerable stability and confidence in the best cluster of docked complexes.

RMSD of the antibody/PD-1 complexes (aligned before and after energy minimization in molecular dynamics simulations):

- TUPPD1-001: 1.034Å
- TUPPD1-002: 0.970Å
- TUPPD1-009: 1.072Å
- Pembrolizumab: 0.906Å
- Nivolumab: 0.911Å

### Complex Interface

Furthermore, these aforementioned candidates also form numerous polar contacts with PD-1 at residues similar to that of nivolumab, pembrolizumab, and other reference antibodies. GLN75, GLU84, SER87, LEU128 are common polar contacts seen in these complexes, which are consistent with the general reported interfaces of pembrolizumab (36) and nivolumab (37). Interfaces for TUPPD1-001, TUPPD1-002, and TUPPD1-009 are shown in Figure 3.

**Fig. 3.**
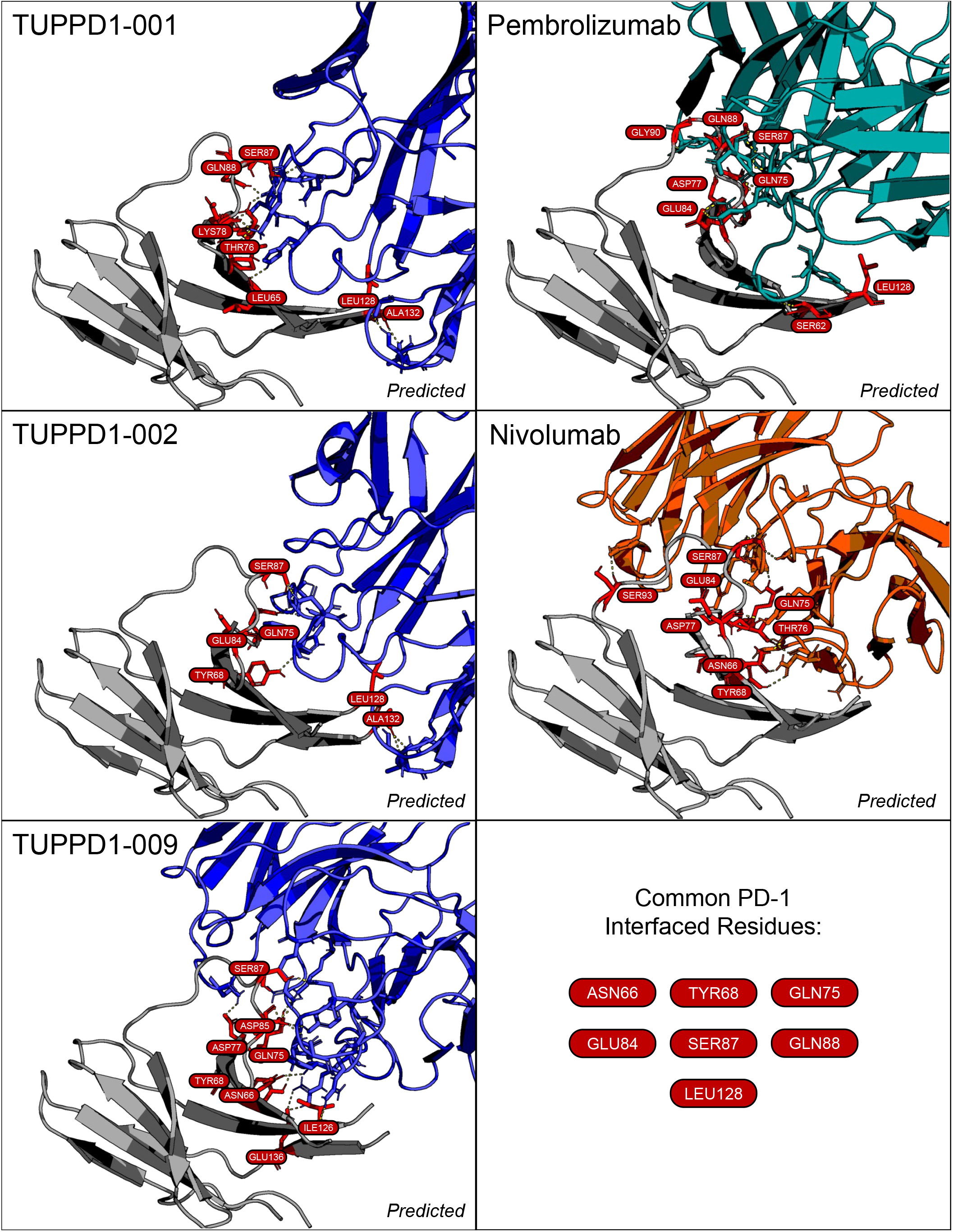
Predicted docking interfaces of three conditionally-diffused antibodies, pembrolizumab, and nivolumab. The grey cartoon protein is PD-1 (with polar contacts shown in red) and the blue cartoon structures are the Fv portions of the diffused antibodies. The Fv structures of pembrolizumab and nivolumab are shown in teal and orange, respectively.

## Discussion

Protein diffusion is an exciting and new technology. Built on the progress in large language models (LLMs) for general artificial intelligence, these models have potential to revolutionize the drug development process and reducing the initial lab-based development workload in favor of *in silico* exploration (38, 39). State-of-the-art protein generation models like EvoDiff from Microsoft Research (20) (used in this study), RFdiffusion (40) from the Baker Lab at the University of Washington, and Chroma (41) from Generate:Biomedicines have shown real promise in AI-based drug design. These various models and frameworks offer a variety of method for protein diffusion. For example, EvoDiff conditionally generates sequences that fold into the expected shape (such as antibody chains) through a simple alignment input. Other models, like Chroma generate proteins based on their potential in a given complex (such as an antibody binding to PD-1). In future studies, we will explore the use of various diffusion models, weighing their respective capabilities for a larger set of diffusion-based *in silico* experiments.

Recently, we have seen clinical trials of computationally-derived antibodies such as Aulos Bioscience’s AU-007 antibody targeting IL-2 in patients with unresectable locally advanced or metastatic cancer (42). Thus, the potential of *in silico*-based therapeutics *in vivo* is certainly nascent.

## Conclusion

For anti-PD-1 therapeutics, here we have shown the utility of conditional diffusion in the guidance of amino acid sequence generation that mimics the biophysical features of other available antibodies that target PD-1.

The results presented here are limited in that they are *in silico*-based predictions. Additional testing will be performed in the future to empirically validate these findings. This will include antibody synthesis, testing on recombinant cell lines, and other lab-based assessments of binding affinity.

Furthermore, the diffusion process presented here only includes the creation of the Fv portion of the candidate structures. Thus, additional work is left to be performed to complete the full antibody structure, including the combination with the rest of the immunoglobulin G structure, any humanization or efforts to reduce immunogenicity, etc. Since it appears that some of the biochemical features of these diffused sequences may be unusual as compared to naturally-derived antibodies, it is still unknown how this will affect the viability of the candidates, both from a therapeutic perspective and from a manufacturability perspective. For example, from the abYsis analyses, we see that various residues are uncommon in certain positions, but we do not yet understand how those residues may impact the an antibody’s potential (positively or negatively) to be developed into a safe and effective drug.

Previous studies by our team (30, 43–45) and others (46–48) have shown the utility of large-scale computational screens for understanding the interaction of antibodies (or other immunoproteins) with protein targets as well as the identification of therapeutic targets of interest. Combining this capability with protein diffusion, we can increase the throughput of computational drug design through high-performance computing and automated complex evaluation, as we’ve demonstrated in this preliminary work.

## Data availability statement

All code, data, results, and additional analyses are openly available on GitHub at: https://github.com/tuplexyz/PD-1_Fab_Diffusion.

This repository includes the PDB files for the all 42 antibody Fv structures (33 from Thera-SAbDab and the 9 diffusion-based structures), the Fv-PD-1 complexes from HADDOCK, all output metrics, experiment generation and data preparation logic, HPC submission scripts, and the code for post-processing analyses and generating figures.

## Disclosures

### Trademarks

Keytruda® (pembrolizumab) is a registered trademark of Merck Sharp & Dohme Corp.

Opdivo® (nivolumab) is a registered trademark of Bristol-Myers Squibb Company.

AU-007 is a clinical antibody candidate owned by Aulos Bioscience.

## Footnotes

1 Container GitHub Repository: https://github.com/colbyford/HADDOCKer

2 Docker Hub Images: https://hub.docker.com/r/cford38/haddock

## References

1. David S. Wilkes and William J. Burlingham. Immunobiology of Organ Transplantation. Springer, 2012. ISBN 9781441989994.

2. Sara Gandini, Daniela Massi, and Mario Mandalà. PD-L1 expression in cancer patients receiving anti PD-1/PD-L1 antibodies: A systematic review and meta-analysis. Critical Reviews in Oncology/Hematology, 100:88–98, 2016. ISSN 1040-8428. doi: 10.1016/j.critrevonc.2016.02.001.

3. Jinming Yu, Xin Wang, Feifei Teng, and Li Kong. PD-L1 expression in human cancers and its association with clinical outcomes. OncoTargets and Therapy, Volume 9:5023–5039, 2016. doi:10.2147/ott.s105862.

4. Hashem O. Alsaab, Samaresh Sau, Rami Alzhrani, Katyayani Tatiparti, Ketki Bhise, Sushil K. Kashaw, and Arun K. Iyer. PD-1 and PD-L1 Checkpoint Signaling Inhibition for Cancer Immunotherapy: Mechanism, Combinations, and Clinical Outcome. Frontiers in Pharmacology, 8, 2017. ISSN 1663-9812. doi:10.3389/fphar.2017.00561.

5. Drug Approval Package - Keytruda (pembrolizumab) Powder for Injection, Oct 2014.

6. Drug Approval Package - Opdivo (nivolumab) Injection, Dec 2014.

7. Anasuya Gunturi and David F McDermott. Nivolumab for the treatment of cancer. Expert Opinion on Investigational Drugs, 24(2):253–260, 2015. doi:10.1517/13543784.2015.991819. PMID: 25494679.

8. Raghav Sundar, Byoung-Chul Cho, Julie R. Brahmer, and Ross A. Soo. Nivolumab in nsclc: latest evidence and clinical potential. Therapeutic Advances in Medical Oncology, 7(2):85–96, 2015. doi:10.1177/1758834014567470. PMID: 25755681.

9. Douglas B. Johnson, Chengwei Peng, and Jeffrey A. Sosman. Nivolumab in melanoma: latest evidence and clinical potential. Therapeutic Advances in Medical Oncology, 7(2):97–106, 2015. doi:10.1177/1758834014567469. PMID: 25755682.

10. Robert J. Motzer, Bernard Escudier, David F. McDermott, Saby George, Hans J. Hammers, Sandhya Srinivas, Scott S. Tykodi, Jeffrey A. Sosman, Giuseppe Procopio, Elizabeth R. Plimack, Daniel Castellano, Toni K. Choueiri, Howard Gurney, Frede Donskov, Petri Bono, John Wagstaff, Thomas C. Gauler, Takeshi Ueda, Yoshihiko Tomita, Fabio A. Schutz, Christian Kollmannsberger, James Larkin, Alain Ravaud, Jason S. Simon, Li-An Xu, Ian M. Waxman, and Padmanee Sharma. Nivolumab versus Everolimus in Advanced Renal-Cell Carcinoma. New England Journal of Medicine, 373(19):1803–1813, 2015. doi:10.1056/NEJMoa1510665. PMID: 26406148.

11. Kortnye Maureen Smith and Jayesh Desai. Nivolumab for the treatment of colorectal cancer. Expert Review of Anticancer Therapy, 18(7):611–618, 2018. doi:10.1080/14737140.2018.1480942. PMID: 29792730.

12. Naiyer A. Rizvi, Matthew D. Hellmann, Alexandra Snyder, Pia Kvistborg, Vladimir Makarov, Jonathan J. Havel, William Lee, Jianda Yuan, Phillip Wong, Teresa S. Ho, Martin L. Miller, Natasha Rekhtman, Andre L. Moreira, Fawzia Ibrahim, Cameron Bruggeman, Billel Gasmi, Roberta Zappasodi, Yuka Maeda, Chris Sander, Edward B. Garon, Taha Merghoub, Jedd D. Wolchok, Ton N. Schumacher, and Timothy A. Chan. Mutational landscape determines sensitivity to PD-1 blockade in non–small cell lung cancer. Science, 348(6230):124–128, 2015. doi:10.1126/science.aaa1348.

13. Gerry Kwok, Thomas C. C. Yau, Joanne W. Chiu, Eric Tse, and Yok-Lam Kwong. Pembrolizumab (Keytruda). Human Vaccines & Immunotherapeutics, 12(11):2777–2789, 2016. doi:10.1080/21645515.2016.1199310. PMID: 27398650.

14. Nicholas L. Syn, Michele W. L. Teng, Tony S. K. Mok, and Ross A. Soo. De-novo and acquired resistance to immune checkpoint targeting. The Lancet Oncology, 18(12):e731– e741, Dec 2017. ISSN 1470-2045. doi:10.1016/S1470-2045(17)30607-1.

15. Saman Maleki Vareki, Carmen Garrigós, and Ignacio Duran. Biomarkers of response to PD-1/PD-L1 inhibition. Critical Reviews in Oncology/Hematology, 116:116–124, 2017. ISSN 1040-8428. doi: 10.1016/j.critrevonc.2017.06.001.

16. Hanxiao Li, P. Anton van der Merwe, and Shivan Sivakumar. Biomarkers of response to PD-1 pathway blockade. British Journal of Cancer, 126(12):1663–1675, Jun 2022. ISSN 1532-1827. doi:10.1038/s41416-022-01743-4.

17. Youhai Jiang, Xiaofang Zhao, Jing Fu, and Hongyang Wang. Progress and Challenges in Precise Treatment of Tumors With PD-1/PD-L1 Blockade. Frontiers in Immunology, 11, 2020. ISSN 1664-3224. doi:10.3389/fimmu.2020.00339.

18. Matthew I J Raybould, Claire Marks, Alan P Lewis, Jiye Shi, Alexander Bujotzek, Bruck Taddese, and Charlotte M Deane. Thera-SAbDab: the Therapeutic Structural Antibody Database. Nucleic Acids Research, 48(D1):D383–D388, 09 2019. ISSN 0305-1048. doi:10.1093/nar/gkz827.

19. Robert C. Edgar. MUSCLE: a multiple sequence alignment method with reduced time and space complexity. BMC Bioinformatics, 5(1):113, Aug 2004. ISSN 1471-2105. doi:10.1186/1471-2105-5-113.

20. Sarah Alamdari, Nitya Thakkar, Rianne van den Berg, Alex X. Lu, Nicolo Fusi, Ava P. Amini, and Kevin K. Yang. Protein generation with evolutionary diffusion: sequence is all you need. bioRxiv, 2023. doi:10.1101/2023.09.11.556673.

21. Mark B. Swindells, Craig T. Porter, Matthew Couch, Jacob Hurst, K.R. Abhinandan, Jens H. Nielsen, Gary Macindoe, James Hetherington, and Andrew C.R. Martin. abYsis: Integrated Antibody Sequence and Structure—Management, Analysis, and Prediction. Journal of Molecular Biology, 429(3):356–364, 2017. ISSN 0022-2836. doi: 10.1016/j.jmb.2016.08.019. Computation Resources for Molecular Biology.

22. Milot Mirdita, Konstantin Schütze, Yoshitaka Moriwaki, Lim Heo, Sergey Ovchinnikov, and Martin Steinegger. ColabFold: making protein folding accessible to all. Nature Methods, 19 (6):679–682, Jun 2022. ISSN 1548-7105. doi:10.1038/s41592-022-01488-1.

23. John Jumper, Richard Evans, Alexander Pritzel, Tim Green, Michael Figurnov, Olaf Ronneberger, Kathryn Tunyasuvunakool, Russ Bates, Augustin Žídek, Anna Potapenko, Alex Bridgland, Clemens Meyer, Simon A. A. Kohl, Andrew J. Ballard, Andrew Cowie, Bernardino Romera-Paredes, Stanislav Nikolov, Rishub Jain, Jonas Adler, Trevor Back, Stig Petersen, David Reiman, Ellen Clancy, Michal Zielinski, Martin Steinegger, Michalina Pacholska, Tamas Berghammer, Sebastian Bodenstein, David Silver, Oriol Vinyals, Andrew W. Senior, Koray Kavukcuoglu, Pushmeet Kohli, and Demis Hassabis. Highly accurate protein structure prediction with AlphaFold. Nature, 596(7873):583–589, Aug 2021. ISSN 1476-4687. doi:10.1038/s41586-021-03819-2.

24. Richard Evans, Michael O’Neill, Alexander Pritzel, Natasha Antropova, Andrew Senior, Tim Green, Augustin Žídek, Russ Bates, Sam Blackwell, Jason Yim, Olaf Ronneberger, Sebastian Bodenstein, Michal Zielinski, Alex Bridgland, Anna Potapenko, Andrew Cowie, Kathryn Tunyasuvunakool, Rishub Jain, Ellen Clancy, Pushmeet Kohli, John Jumper, and Demis Hassabis. Protein complex prediction with AlphaFold-Multimer. bioRxiv, 2022. doi:10.1101/2021.10.04.463034.

25. Martin Steinegger and Johannes Söding. MMseqs2 enables sensitive protein sequence searching for the analysis of massive data sets. Nature Biotechnology, 35(11):1026–1028, Nov 2017. ISSN 1546-1696. doi:10.1038/nbt.3988.

26. Peter Eastman, Jason Swails, John D. Chodera, Robert T. McGibbon, Yutong Zhao, Kyle A. Beauchamp, Lee-Ping Wang, Andrew C. Simmonett, Matthew P. Harrigan, Chaya D. Stern, Rafal P. Wiewiora, Bernard R. Brooks, and Vijay S. Pande. Openmm 7: Rapid development of high performance algorithms for molecular dynamics. PLOS Computational Biology, 13 (7):1–17, 07 2017. doi:10.1371/journal.pcbi.1005659.

27. Schrödinger, LLC. The PyMOL molecular graphics system, version 1.8. November 2015.

28. G.C.P. van Zundert, J.P.G.L.M. Rodrigues, M. Trellet, C. Schmitz, P.L. Kastritis, E. Karaca, A.S.J. Melquiond, M. van Dijk, S.J. de Vries, and A.M.J.J. Bonvin. The HADDOCK2.2 Web Server: User-Friendly Integrative Modeling of Biomolecular Complexes. Journal of Molecular Biology, 428(4):720–725, 2016. doi:10.1016/j.jmb.2015.09.014.

29. James Dunbar and Charlotte M. Deane. ANARCI: antigen receptor numbering and receptor classification. Bioinformatics, 32(2):298–300, 09 2015. ISSN 1367-4803. doi:10.1093/bioinformatics/btv552.

30. Phillip J. Tomezsko, Colby T. Ford, Avery E. Meyer, Adam M. Michaleas, and Rafael Jaimes. Human cytokine and coronavirus nucleocapsid protein interactivity using large-scale virtual screens. Frontiers in Bioinformatics, 4, 2024. ISSN 2673-7647. doi:10.3389/fbinf.2024.1397968.

31. Francesco Ambrosetti, Brian Jiménez-García, Jorge Roel-Touris, and Alexandre M.J.J. Bonvin. Modeling antibody-antigen complexes by information-driven docking. Structure, 28(1):119–129.e2, 2020. ISSN 0969-2126. doi: 10.1016/j.str.2019.10.011.

32. Francesco Ambrosetti, Zuzana Jandova, and Alexandre M. J. J. Bonvin. “Information-Driven Antibody–Antigen Modelling with HADDOCK”, pages 267–282. Springer US, New York, NY, 2023. ISBN 978-1-0716-2609-2. doi:10.1007/978-1-0716-2609-2_14.

33. Anna Vangone and Alexandre MJJ Bonvin. Contacts-based prediction of binding affinity in protein–protein complexes. eLife, 4:e07454, jul 2015. ISSN 2050-084X. doi:10.7554/eLife.07454.

34. Jumin Lee, Xi Cheng, Jason M. Swails, Min Sun Yeom, Peter K. Eastman, Justin A. Lemkul, Shuai Wei, Joshua Buckner, Jong Cheol Jeong, Yifei Qi, Sunhwan Jo, Vijay S. Pande, David A. Case, Charles L. Brooks III, Alexander D. MacKerell Jr., Jeffery B. Klauda, and Wonpil Im. CHARMM-GUI Input Generator for NAMD, GROMACS, AMBER, OpenMM, and CHARMM/OpenMM Simulations Using the CHARMM36 Additive Force Field. Journal of Chemical Theory and Computation, 12(1):405–413, Jan 2016. ISSN 1549-9618. doi:10.1021/acs.jctc.5b00935.

35. Tom Darden, Darrin York, and Lee Pedersen. Particle mesh Ewald: An Nlog(N) method for Ewald sums in large systems. The Journal of Chemical Physics, 98(12):10089–10092, 06 1993. ISSN 0021-9606. doi:10.1063/1.464397.

36. Zhenkun Na, Siok Ping Yeo, Sakshibeedu R. Bharath, Matthew W. Bowler, Esra Balıkçı, Cheng-I Wang, and Haiwei Song. Structural basis for blocking PD-1-mediated immune suppression by therapeutic antibody pembrolizumab. Cell Research, 27(1):147–150, Jan 2017. ISSN 1748-7838. doi:10.1038/cr.2016.77.

37. Shuguang Tan, Hao Zhang, Yan Chai, Hao Song, Zhou Tong, Qihui Wang, Jianxun Qi, Gary Wong, Xiaodong Zhu, William J. Liu, Shan Gao, Zhongfu Wang, Yi Shi, Fuquan Yang, George F. Gao, and Jinghua Yan. An unexpected N-terminal loop in PD-1 dominates binding by nivolumab. Nature Communications, 8(1):14369, Feb 2017. ISSN 2041-1723. doi:10.1038/ncomms14369.

38. Ganggang Bai, Chuance Sun, Ziang Guo, Yangjing Wang, Xincheng Zeng, Yuhong Su, Qi Zhao, and Buyong Ma. Accelerating antibody discovery and design with artificial intelligence: Recent advances and prospects. Seminars in Cancer Biology, 95:13–24, 2023. ISSN 1044-579X. doi: 10.1016/j.semcancer.2023.06.005.

39. Ge Liu, Haoyang Zeng, Jonas Mueller, Brandon Carter, Ziheng Wang, Jonas Schilz, Geraldine Horny, Michael E Birnbaum, Stefan Ewert, and David K Gifford. Antibody complementarity determining region design using high-capacity machine learning. Bioinformatics, 36 (7):2126–2133, 11 2019. ISSN 1367-4803. doi:10.1093/bioinformatics/btz895.

40. Joseph L. Watson, David Juergens, Nathaniel R. Bennett, Brian L. Trippe, Jason Yim, Helen E. Eisenach, Woody Ahern, Andrew J. Borst, Robert J. Ragotte, Lukas F. Milles, Basile I. M. Wicky, Nikita Hanikel, Samuel J. Pellock, Alexis Courbet, William Sheffler, Jue Wang, Preetham Venkatesh, Isaac Sappington, Susana Vázquez Torres, Anna Lauko, Valentin De Bortoli, Emile Mathieu, Sergey Ovchinnikov, Regina Barzilay, Tommi S. Jaakkola, Frank DiMaio, Minkyung Baek, and David Baker. De novo design of protein structure and function with RFdiffusion. Nature, 620(7976):1089–1100, Aug 2023. ISSN 1476-4687. doi:10.1038/s41586-023-06415-8.

41. John B. Ingraham, Max Baranov, Zak Costello, Karl W. Barber, Wujie Wang, Ahmed Ismail, Vincent Frappier, Dana M. Lord, Christopher Ng-Thow-Hing, Erik R. Van Vlack, Shan Tie, Vincent Xue, Sarah C. Cowles, Alan Leung, João V. Rodrigues, Claudio L. Morales-Perez, Alex M. Ayoub, Robin Green, Katherine Puentes, Frank Oplinger, Nishant V. Panwar, Fritz Obermeyer, Adam R. Root, Andrew L. Beam, Frank J. Poelwijk, and Gevorg Grigoryan. Illuminating protein space with a programmable generative model. Nature, 623(7989):1070– 1078, Nov 2023. ISSN 1476-4687. doi:10.1038/s41586-023-06728-8.

42. Inc. Aulos Bioscience. A Study of AU-007 in Adult Subjects With Advanced Solid Tumors, 2024. ClinicalTrials.gov Identifier: NCT05267626.

43. Colby T. Ford, Denis Jacob Machado, and Daniel A Janies. Predictions of the SARS-CoV-2 Omicron Variant (B.1.1.529) Spike Protein Receptor-Binding Domain Structure and Neutralizing Antibody Interactions. Frontiers in Virology, 2022. doi:10.3389/fviro.2022.830202.

44. COlby T. Ford, Shirish Yasa, Denis Jacob Machado, Richard Allen White III, and Daniel A. Janies. Predicting changes in neutralizing antibody activity for SARS-CoV-2 XBB.1.5 using in silico protein modeling. Frontiers in Virology, 3, 2023. ISSN 2673-818X. doi:10.3389/fviro.2023.1172027.

45. Cheikh Cambel Dieng, Colby T. Ford, Anita Lerch, Dickson Doniou, Kovidh Vegesna, Daniel Janies, Liwang Cui, Linda Amoah, Yaw Afrane, and Eugenia Lo. Genetic variations of plasmodium falciparum circumsporozoite protein and the impact on interactions with human immunoproteins and malaria vaccine efficacy. Infection, Genetics and Evolution, 110:105418, 2023. ISSN 1567-1348. doi: 10.1016/j.meegid.2023.105418.

46. Marco Giulini, Constantin Schneider, Daniel Cutting, Nikita Desai, Charlotte M. Deane, and Alexandre M.J.J. Bonvin. Towards the accurate modelling of antibody-antigen complexes from sequence using machine learning and information-driven docking. bioRxiv, 2023. doi: 10.1101/2023.11.17.567543.

47. Francis Gaudreault, Christopher R. Corbeil, and Traian Sulea. Enhanced antibody-antigen structure prediction from molecular docking using AlphaFold2. Scientific Reports, 13(1):15107, Sep 2023. ISSN 2045-2322. doi: 10.1038/s41598-023-42090-5.

48. Mohsen M. Seyed, Fateme Sefid, Sahar Shahqoli, and Parisa Sharifi. Antibody engineering to increase the affinity of edrecolomab monoclonal antibody against vegf-a protein in the treatment of colorectal cancer. NeuroQuantology, 21(7):263–279, 2023. Copyright - Copyright NeuroQuantology 2023; Last updated - 2023-12-19.

